# The Impact of BOLD Induced Linewidth Modulation on Functional ^1^H MRS Analysis

**DOI:** 10.64898/2026.03.06.710034

**Authors:** Martin Wilson, Simon M. Finney, William T. Clarke

## Abstract

Functional MRS can measure the neurometabolic response to neuronal activation, therapeutic interventions and changes in physiology. Substantial technical challenges currently present a barrier to reproducible findings and broader adoption by the neuroscientific community. One such challenge is the conflation between genuine metabolic changes and bias caused by subtle spectral lineshape changes associated with the BOLD response. Previous studies have demonstrated an approximately 1% bias for glutamate estimates at 7T based on experimentally acquired data and a single conventional fitting algorithm. In this study, we use synthetic MRS data to estimate the bias for two conventional fitting methods (LCModel and ABfit-reg) at 3T and 7T and evaluate the efficacy of dynamic lineshape adjustment, during preprocessing and fitting analysis stages, to reduce bias. Using the same dataset, we also explore the potential bias in 2D fitting approaches, comparing several fitting models implemented in FSL-MRS. Bias between two conventional fitting methods without explicit linewidth correction was similar (∼1% for glutamate) and in good agreement with previous experimental studies at 7T. Lineshape changes from the BOLD response cause similar bias in conventional and 2D fitting packages for 3T and 7T data, resulting in an overestimation of metabolic changes associated with neuronal activation. This bias may be significantly reduced (<0.2%) by incorporating a BOLD linewidth matching step for conventional analysis or by direct modelling for 2D analysis. We therefore recommend explicit BOLD lineshape correction or modelling for future task-based fMRS studies at 3T and above.

## Introduction

Functional magnetic resonance spectroscopy (fMRS) is a non-invasive contrast that measures dynamic changes in metabolite signal associated with neuronal activation^1,2^. Early studies demonstrated an increase in visual cortex lactate in response to a prolonged period (∼10 minutes) of visual stimulation^3^, and these findings have been replicated by various research groups across multiple acquisition platforms and analysis methodologies^4–11^. More recently, methodological advances have established associated changes in glutamate, glucose and aspartate^4,6,9,10^, providing novel information on the roles of the aerobic glycolytic pathway, malate–aspartate shuttle and tricarboxylic acid cycle in neural activity. Alternative paradigms, including pain^12–14^, motor tasks^15–17^ and event-related designs^18,19^ have also demonstrated various changes in metabolite levels in response to brain activation.

Whilst fMRS provides novel molecular insight into localised brain activation^20^, the technique is technically challenging to perform. This is primarily due to the relatively low signal strength of the metabolites compared to spectral confounds, such as thermal noise, and the insufficient suppression of water and lipid signals. The positive BOLD effect presents a further counter-intuitive confound by inducing a small reduction in spectral linewidth during periods of increased brain activation^21,22^. In principle, established spectral fitting methods^23^ model lineshape changes, implying they should not influence metabolite levels. However, comparisons of fitting models have found biased estimations of lineshape in some packages^24^ and fMRS studies at 7T have demonstrated how minor spectral narrowing biases dynamic metabolite estimates^4,25^. This bias presents a significant concern for fMRS interpretation, as genuine task induced changes in metabolite levels are generally concurrent with the BOLD effect, and therefore BOLD induced fitting bias may be erroneously attributed to genuine changes in metabolite levels.

At the time of writing, the fMRS community is yet to reach consensus on precisely which experimental conditions warrant explicit correction of the BOLD induced bias, with prevailing opinion suggesting these effects only need to be ameliorated at field strengths of 7T and above. With this opinion presumably being motivated by the BOLD effects linear dependence on field strength^26^. The most widely used correction involves applying Lorenzian linebroadening to the spectra acquired during the task-active portion of the experiment to match those acquired in the rest or baseline phase^4,10,17,27^. We will refer to this approach as “Lorentzian lineshape matching” in this paper. Additionally, white noise may be added to the artificially broadened spectra to eliminate any potential SNR bias between the rest and task phases^28^. Both these steps are performed during the preprocessing phase of analysis before spectral fitting^29^.

In this study, we quantify the magnitude of the BOLD induced bias on metabolite estimates and assess several strategies to mitigate the effect. In contrast to previous studies, based on in-vivo data^4,25^, we use synthetic MRS to precisely isolate the BOLD induced bias from other potential confounds – such as participant movement or scanner drift^30^. Furthermore, our analysis goes beyond conventional 1D fitting at 7T to also include 3T and 2D dynamic fitting^31–33^, covering a wider range of contemporary fMRS methodologies. We also address the effect of noise colouring introducing bias during fitting when the Lorentzian lineshape matching approach, also known as apodisation, is used.

## Methods

The generation and analysis of synthetic MRS is currently the only approach to offer “ground-truth” knowledge of the dynamic spectral changes associated with fMRS. Therefore, our methodology uses simulation of the expected dynamic lineshape change associated with the BOLD response to a prolonged task - currently the most reproduced fMRS paradigm. Simulation parameters, such as SNR, linewidth and metabolite levels, are designed to match those from previously published studies acquired at field strengths of 3T and 7T. In contrast to experimentally acquired MRS, simulated metabolite levels will not change dynamically, therefore any task-associated change in metabolite levels may be directly interpreted as BOLD induced bias, i.e. the expected ground truth result is for no metabolite concentration change.

### Synthetic data

The list of metabolite, macromolecule and lipid signals and concentrations used to generate the synthetic MRS data is given in Table S1. Metabolite chemical shift and J-coupling values were taken from Govindaraju et al^34,35^, broad lipid and macromolecule signal parameters were taken from Table 1 in Wilson et al^36^. Metabolite concentration values, emulating visual cortex, were taken from Table 1 in Bednarik et al^4^. Alanine and Glycine were not listed in this table and therefore did not contribute to the synthetic MRS data, however both metabolites were included in the subsequent fitting basis set.

**Table 1.** Simulation parameters for 3 T and 7 T synthetic datasets.

|  | 3 T | 7 T |
| --- | --- | --- |
| **Single shot SNR** | 45 | 80 |
| **TR (seconds)** | 2.5 | 5 |
| **Number of excitations** | 640 | 320 |
| **Applied Gaussian broadening (Hz)** | 6 | 11 |
| **BOLD Lorentzian linewidth change (Hz)** | 0.2 | 0.46 |

Metabolite signals were simulated for the semi-LASER sequence^37^ with an echo time of 28 ms, acquired over 2048 complex points sampled at a frequency of 6000 Hz. The total scan time was 26 minutes and 40 seconds. Table 1 lists simulation parameters that differ between 3 T and 7 T based on typical values for visual cortex acquisitions reported in the literature^4^. Example spectra are plotted in Figure S1.

A simple prolonged block design was assumed to generate the BOLD linewidth changes. Five blocks of equal duration (320 seconds) were ordered as REST-TASK-REST-TAST-REST, with the TASK blocks corresponding to a reduction in Lorentzian linewidth of 0.2 Hz and 0.46 Hz at 3 T and 7 T respectively, based on previous studies of fMRS visual stimulation^4,22^. In practice, Lorentzian linebroadening was applied to the REST blocks and its contribution reduced in the TASK regions assuming a BOLD response derived from the widely used double gamma model^38^. Figure 1 shows the applied Lorentizain linebroadening for both field strengths, with the notable feature that a constant value was added to both responses to ensure all values were non-negative to avoid numerical instability.

**Figure 1.**
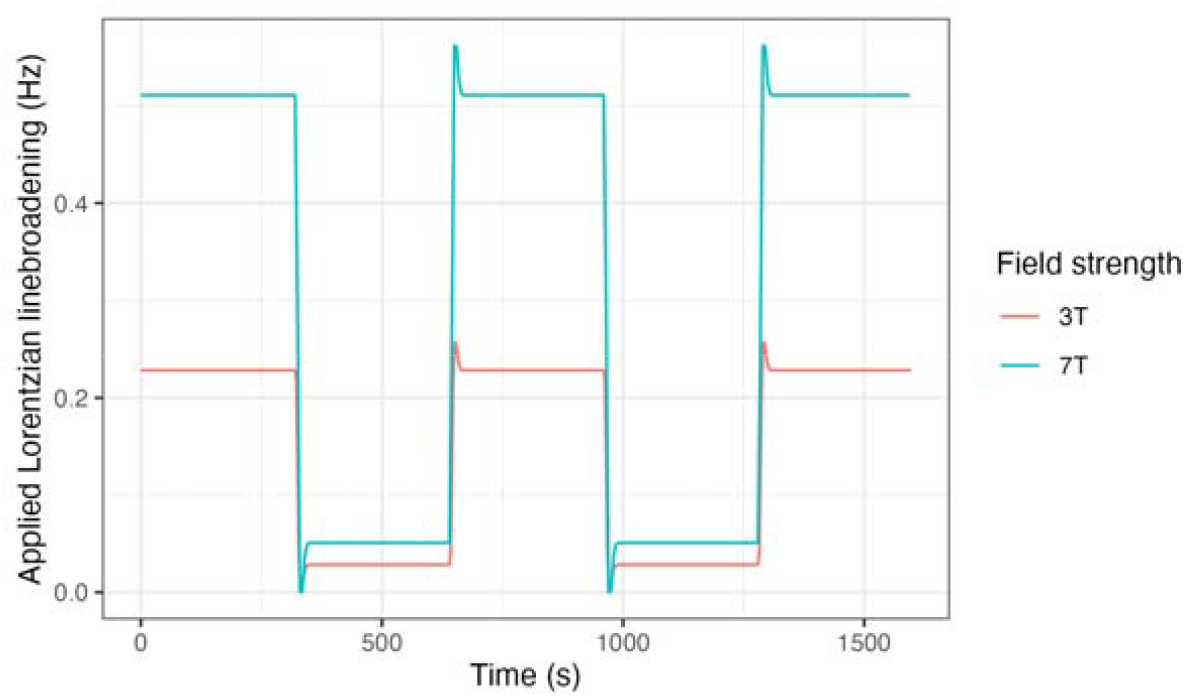
Applied Lorentizan linebroadening to simulate the spectral BOLD effect at 3 T and 7 T.

The full dataset consisted of 128 synthetic fMRS scans with additive Gaussian distributed complex noise to match the single-shot SNR values given in Table 1. This number of scans was chosen to be large enough to accurately estimate relevant bias from typical fMRS experiments, which generally consist of less than 20 participants. Each fMRS scan was designated a subject ID (eg sub-001, sub-002…) stored in NIfTI-MRS format^39^ and organised according to BIDS specification^40^.

### Lorentzian lineshape matching

The most common method to reduce BOLD line-narrowing induced fitting bias implements a data preprocessing step where Lorentzian linebroadening is applied to data acquired during the TASK phase of the experiment to match the REST phase. In this study, we have also applied the Lorentzian lineshape matching preprocessing step to the synthetic data to evaluate its efficacy in reducing fitting bias. In addition to applying Lorentzian linebroadening to maintain linewidth consistency across the dynamic scans (based on the known curves shown in Figure 1), we also add additional complex Gaussian noise to maintain SNR consistency – as performed by Bednarik at al^4^. Figure S2 illustrates this procedure, with comparison between parts B and C showing the lineshape matching step and C and D showing the noise matching step. Analysis incorporating this approach will be denoted “preproc lw matching” in subsequent plots.

### Conventional fitting and preprocessing

Before fitting, Lorentzian lineshape matching (as described above) was applied to a copy of the data for the “preproc lw matching” analysis, whilst the original un-matched was used for all other analyses (“basis lw matching” and “default analysis”). Dynamic frequency and phase alignment is a standard preprocessing step for fMRS, therefore the RATS method^41^ was applied to each dataset. Averaging of temporal dynamics was also applied to reduce the original, unaveraged data down to 5 blocks (REST, TASK, REST, TASK, REST). Two alternative averaging schemes, 2 blocks (REST, TASK), and 10 blocks (REST, REST, TASK, TASK, REST, REST, TASK, TASK, REST, REST) were also assessed.

Two fitting packages were used:

1. LCModel^23^, as this package has been reported to exhibit BOLD induced bias^25^ and is widely used for fMRS analysis,
2. ABfit-reg^42^, to explore if bias differs significantly between conventional fitting algorithms.

Default fitting options were used for both fitting packages, and the same basis set used for synthetic MRS generation was used for spectral fitting (Table S1).

As an alternative to the conventional preprocessing lineshape matching step, we performed an additional analysis where the linewidth of the fitting basis set was adjusted (prior to subsequent adjustment within the fitting method) to compensate for the change arising from the predicted BOLD response. For example, in the 3 T data analysis, 0.2 Hz Lorentzian linebroadening was applied to the basis set for the REST blocks, but not the TASK blocks. This was performed as an attempt to equalise the magnitude of any initial fitting discrepancy (and therefore potential bias) between the REST and TASK states. Analysis incorporating this approach is denoted “basis lw matching” in subsequent results.

Synthetic MRS generation, preprocessing and conventional fitting were performed using version of the spant MRS analysis package^43^.

### Dynamic fitting

Dynamic fitting, the simultaneous modelling of the spectral and (functional) temporal domains, was also assessed as a method to minimise line-narrowing bias. This assessment was carried out using FSL-MRS ^31,44^ version 2.4.10, on the same data (simulated as described above), and using the same fitting basis sets. Three different temporal models were investigated, in each the time-dependance of metabolite concentrations were modeled using a GLM with three regressors: a “constant” term, and two rectangular functions aligned with the periods of BOLD-induced line-narrowing. The models differed in their treatment of the time-dependance of Lorentzian linebroadening term (the FSL-MRS “gamma” parameter):

1. Fixed Model The Fixed Model imposed a constant, but optimisable value of Lorentzian broadening for all timepoints.
2. Exact Model The Exact Model imposes the true time-variable broadening used to simulate the data (Figure 1), but the amplitude of the broadening is estimated directly from the data.
3. Correct Model The Correct Model imposes the true time-variable broadening at a scale set to that used to simulate the data.

For the Exact Model, the single linewidth scaling parameter was initialised using a Tobit (censored) regression to mitigate the effect of zero-valued Lorentzian linewidth estimates from single noisy transients. All other model parameters (frequency shift, zero-order and first-order phase, gaussian line broadening) were fitted with a single, optimisable, but time-invariant value. The FSL-MRS baseline was disabled to match the simulated data model. The three dynamic models are provided in the linked Github repository https://github.com/wtclarke/fmrs_bold_dynamic_fitting in the dyn_models directory (dyn_models/config_correct_lb.py, dyn_models/config_exact_lb.py, dyn_models/config_fixed_lb.py)

The FSL tool FLAMEO, which implements multilevel linear modeling for group analysis using Bayesian inference^45^, was used through FSL-MRS’s *fmrs_stats* tool to calculate single-group average statistics for all metabolites. The two blocks of line-narrowing were combined as an equally weighted first-level contrast. The group variance was modelled with fixed effects (runmode=fe) as the simulated data includes no inter-subject variance. The results were calculated as the average estimated change in concentration during the two line-narrowing periods. The FLAMEO estimated group effect, the standard deviation, and Cohen’s d are reported for major metabolites. Note all standard deviations (conventional and dynamic) are calculated as the standard deviation of the per-subject bias, i.e., for the dynamic fitting the reported standard deviation is calculated on the first-level (per-subject) bias.

### Bias in fitting apodised data

Lorentzian lineshape matching, the most widely implemented correction in the literature (and first addressed in this work), applies Lorenzian linebroadening, equivalent to an apodisation filter, to the time-domain FID. The application of this filter affects both the signal and noise in the FID. The noise in the unfiltered FID and the equivalent spectrum (after Fourier transform) is complex, zero-mean, independent and identically distributed (i.i.d.) Gaussian. In the spectral domain, application of the time domain filter, will correlate noise in adjacent spectral bins. For a known filter, the form of this correlation can be derived exactly, and is done so for the exponential apodisation (linebroadening) filter in Appendix A1.

MRS fitting software typically implements an ordinary least-squares (OLS) optimisation, which assumes i.i.d. noise. As such the noise correlation introduced by apodisation could bias fitting results. Optimisation methods such as generalised least squares (GLS) may be used to account for the known noise correlation. To explore mitigation of apodisation-induced fitting bias, we use an asymptotic framework to predict the leading-order bias and variance using both OLS and GLS for an idealised fitting problem - fitting a single, lorentzian (FWHM = 5 Hz), on-resonance peak. Details of the asymptotic framework are presented in Appendix A1. A validation of the asymptotic solutions via comparison to Monte Carlo simulations is presented in the supplementary materials (see Figures S3-S5).

## Results

### Conventional analysis

Figure 2A shows the distribution of bias across a number of commonly measured metabolites with, and without, explicit BOLD lineshape modelling using the ABfit-reg fitting method. The default analysis bias ranged between approximately -1% to 5%, with glucose being the metabolite most susceptible to bias at 3T and 7T. Glutamate and lactate are the metabolites most commonly associated with change in fMRS and these exhibited biases of around 1% and 2% respectively for the default analysis. Preprocessing and basis based BOLD linewidth matching approaches both strongly reduced bias for each metabolite by a similar amount. Standard deviations in the bias are shown in Figure 2B, with the preprocessing based BOLD linewidth approach showing a slight increase over the basis and default approaches. This increase in variance is likely due to the artificially added noise samples used to match the spectral SNR between the REST and TASK conditions.

**Figure 2.**
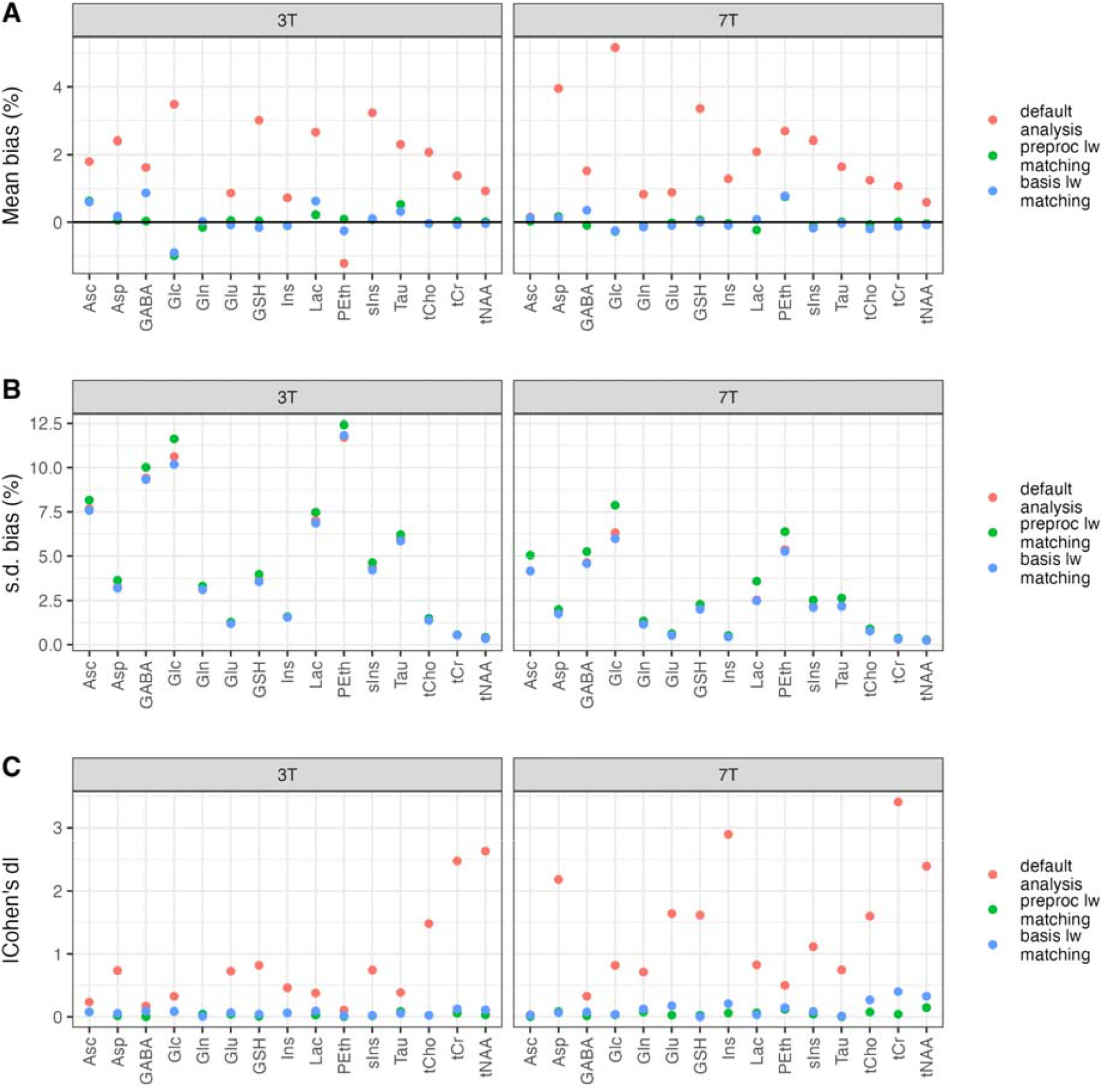
BOLD lineshape induced bias for commonly measured metabolites at 3T and 7T. The ABfit-reg analysis method was used to analyse 5 dynamic blocks (REST - TASK - REST - TASK - REST) and three analysis approaches were compared. Bias was calculated by evaluating the mean of the TASK blocks as a percentage change from the mean of the REST blocks on a participant by participant basis.

An equivalent analysis to Figure 2 for the LCModel fitting method is shown in Figure S7 for comparison with ABfit-reg. In general, the agreement between the two methods is very high, with the biggest differences in biases being restricted to the more poorly determined metabolites such as phosphoethanolamine. Furthermore, the standard deviation of biases (Figures 2B vs S7B) shows a very similar pattern across metabolites, with ABfit-reg showing a small but consistent reduction in standard deviation of around 2.5% for each metabolite.

A more detailed comparison between ABfit-reg and LCModel is shown in Figure S8 for glutamate, with both approaches showing strong agreement. Comparing different dynamic block sizes (2, 5 and 10) shows similar median levels of BOLD lineshape bias between 3T and 7T, however a clear increase in variance is apparent for the 3T results - particularly for the results derived from 10 blocks. This increased variance at 3T indicates that the bias may be less reliably detected for smaller sized studies (N=20) due to metabolite estimation errors being dominated by low SNR rather than linewidth bias.

Table 2 lists the percentage bias for glutamate across the different field strengths and analysis methods. Default analyses gave a bias of around 1% for both LCModel and ABfit-reg independent of field strength. Both preprocessing and basis based linewidth matching approaches reduced the bias to less than 0.09%, which is small enough to be insignificant for typical fMRS designs glutamate is typically expected to change by at least 4%^1^.

**Table 2.** A summary of glutamate bias for the conventional fitting methods, linewidth matching approaches and field strengths investigated. Data were preprocessed into 5 dynamic blocks before fitting.

| Fit method | BOLD linewidth matching approach | Field strength | Mean Glu % change (\|Cohen's d\|) |
| --- | --- | --- | --- |
| LCModel | none | 3 T | 1.00 (0.85) |
| LCModel | none | 7 T | 1.03 (1.83) |
| LCModel | preprocessing | 3 T | 0.06 (0.05) |
| LCModel | preprocessing | 7 T | 0.00 (0.00) |
| LCModel | basis | 3 T | -0.03 (0.02) |
| LCModel | basis | 7 T | 0.05 (0.09) |
| ABfit-reg | none | 3 T | 0.87 (0.73) |
| ABfit-reg | none | 7 T | 0.89 (1.64) |
| ABfit-reg | preprocessing | 3 T | 0.05 (0.04) |
| ABfit-reg | preprocessing | 7 T | -0.02 (0.03) |
| ABfit-reg | basis | 3 T | -0.08 (0.07) |
| ABfit-reg | basis | 7 T | -0.09 (0.18) |

### Dynamic fitting

The results of the dynamic fitting are presented as effect sizes (percentage relative to estimated baseline), standard deviation, and Cohen’s d in Table 3 and Figure 3.

**Table 3.** A summary of glutamate bias for the dynamic fitting methods and field strengths investigated.

| Fit method | Dynamic Fit Model | Field strength | Mean Glu % change (\|Cohen's d\|) |
| --- | --- | --- | --- |
| FSL-MRS | fixed | 3 T | 3.4 (4.3) |
| FSL-MRS | fixed | 7 T | 2.5 (9.5) |
| FSL-MRS | exact | 3 T | -0.6 (0.6) |
| FSL-MRS | exact | 7 T | 0.2 (0.8) |
| FSL-MRS | correct | 3 T | 0.2 (0.2) |
| FSL-MRS | correct | 7 T | 0.1 (0.31) |

**Figure 3.**
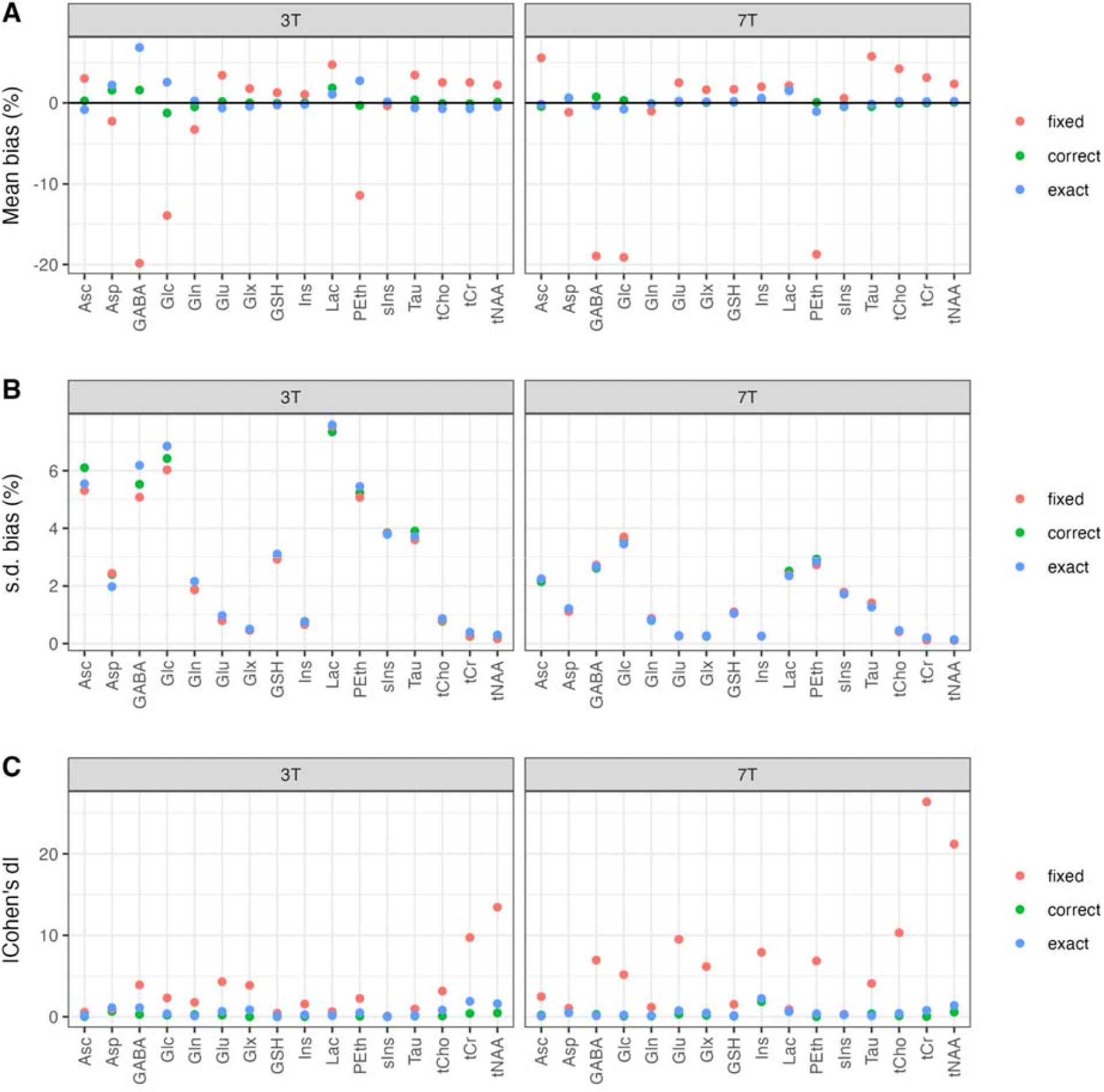
Strip plot of dynamic fitting-measured metabolite concentration changes for each linewidth fitting model during the period of line narrowing. A: Effect size as a percentage of the estimated constant concentration of the metabolite. The horizontal line is the ground truth zero change. B: Standard deviation of the bias across all subjects (expressed as a percentage of the constant concentration). C: Expressed as the magnitude Cohen’s d.

The imposition of an (incorrect) constant linewidth value across all spectra in the *fixed* model results in the largest magnitude biases observed in all models (ranging from -20% to +6%), with glutamate and lactate in the 2% - 5% range at both field strengths. At 3 T 13/16 and at 7T 14/16 metabolites assessed had a Cohen’s d larger than 1.

The *exact* model, which implemented the correct lorentzian lineshape model, but estimated the line narrowing magnitude from the data, showed substantially reduced bias. The overall range was -1% to 7% with Glu showing -0.6 and 0.2 % bias and Lac 1.5 and 1.8 % bias for 3T and 7T respectively. At 3 T 12/16 and at 7T 14/16 metabolites assessed had a Cohen’s d smaller than 1.

The *correct* model fixed the magnitude of the line narrowing to the ground truth value as well as implementing the correct lineshape model. This model provided the lowest bias with the maximum bias, 1.8% shown by Lac at 3T. Glutamate bias was 0.2 (3T) and 0.1 % (7T). At 3 T 16/16 and at 7T 15/16 metabolites assessed had a Cohen’s d smaller than 1, with the average being the lowest of all dynamic models.

### Bias in fitting apodised data

In Figure 4, we present percentage differences between the OLS and GLS predicted concentration bias and standard deviation over a wide range of noise and apodisation magnitudes. The absolute concentration bias and standard deviation for each individual method is presented in Figure S6. In the OLS case, both the bias, but predominantly the variance (and therefore standard deviation) increase with the apodisation. However, using GLS the noise correlation is accounted for and the effect on the bias and variance remains consistent across apodisations, i.e., there is no effect of linebroadening on the concentration. The difference between the OLS and GLS cases therefore informs us to what extent the apodised noise impacts the concentration estimate. We note that for both the OLS and GLS solutions, the primary impact of the noise is to increase the variance of the resulting distribution, since the impact on bias is asymptotically smaller. The concentration standard deviation is 0.01%, 0.03%, and 0.55% larger than unbroadened data for 0.5 Hz, 1 Hz and 10 Hz broadening (OLS, noise = 1.0, equivalent to an unfiltered SNR of 40). Equivalently, the concentration bias is 0.003%, 0.005%, and 0.017% larger than unbroadened data. The effect of apodisation on variance rises substantially with decreasing SNR; with a ten-times lower SNR (of 4), the concentration standard deviation is equivalently 0.11%, 0.32%, and 5.54% larger (than unbroadened data for 0.5 Hz, 1 Hz and 10 Hz broadening).

**Figure 4:**
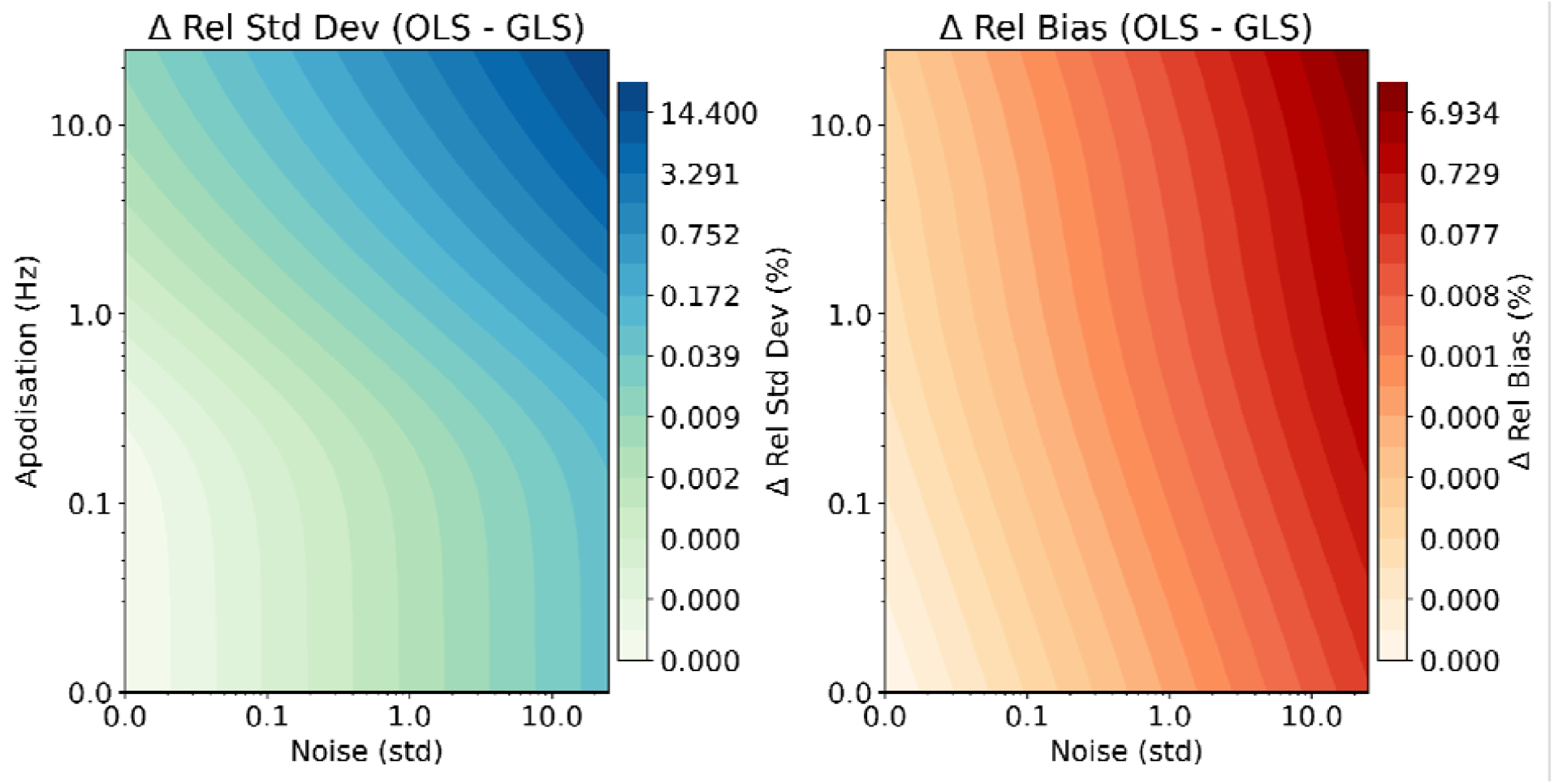
The percentage difference (relative to ground truth) between the fitted concentration standard deviation (left) and bias (right) predicted using the OLS and GLS asymptotic solutions. Results shown over a range of noise and apodisation magnitudes. Contours and axes both use a log scale for visualisation.

## Discussion

Overall, this study justifies the use of explicit BOLD lineshape modelling for conventional and dynamic fMRS analyses of brain activation at 3T and above. Whilst the bias for conventional analyses is low (∼1%) for commonly reported metabolites such as glutamate, a high proportion of studies (15 out of 35) summarised by Mullins et al^1^ reported a glutamate change of less than 5%, suggesting that BOLD bias is significant for certain fMRS studies. Explicit BOLD lineshape modelling is not commonly performed for fMRS at 3T, yet our results show similar bias between 3T and 7T. Whilst the BOLD lineshape change is approximately twice as large (in Hz) at 7T compared to 3T, the reduced spectral overlap and higher SNR at 7T may counteract the increased bias, explaining our observations and strengthening the case for explicit BOLD lineshape modelling at 3T.

Synthetic MRS is an important validation approach for MRS analysis research, since results may be compared directly to the “ground truth”, however oversimplification of the simulation model risks producing results that do not generalise to experimental conditions. Limitations of this work include omitting the effects of subject movement, dynamic baseline instability and dynamic shimming instability. These factors were not considered to ensure any bias could be directly attributed to the BOLD lineshape modulation, and the good agreement between our estimates of glutamate bias and those reported from experimentally acquired fMRS at 7T^4,25^ strongly support our simulation model assumptions.

Lorentzian lineshape matching is the most commonly applied approach to mitigating bias introduced by BOLD. It is achieved by applying a linebroadening apodisation filter to the time-domain signal of the parts of the dataset thought to be affected by BOLD. This work has demonstrated, through an asymptotic framework, that fitting after data apodisation increases both fitting bias and standard deviation. At the level of linebroadening needed to mitigate BOLD at 3 or 7 tesla, i.e., less than 1 hertz, the additional bias and standard deviation introduced is negligible (both <0.1%). With greater BOLD effect at higher proposed field strengths, which at the time of writing include 14 tesla, requiring greater apodisation, or with lower SNR data, arising from the smaller voxels of MRSI, the effect of might not be negligible. The same asymptotic framework was used to validate that the use of GLS fitting removes the effect of apodisation on the fitting. GLS may potentially also remove the need to add additional noise as proposed in reference^4^.

A prolonged visual stimulation task was chosen for the simulated BOLD response due to this being the most replicated fMRS study, however shorter “event-related” stimuli have also been used in fMRS studies^1^. The hemodynamic response function, and therefore BOLD response, is known to reach its maximum value for task durations of around 8 seconds and longer, with event-related designs of 700 ms producing around 15% of this theoretical maximum. Despite this, a clear BOLD response has been observed for event-related fMRS^18^, suggesting it has the potential to introduce fitting bias. However, unlike prolonged tasks, the glutamate response to event-related fMRS is thought to be much faster, peaking 500 ms after stimuli onset^1^, whereas the BOLD response for a 700 ms stimuli would take 5000 ms to reach a maximum. The reduced temporal similarity between the BOLD and glutamate responses for event-related designs should significantly reduce the chances of confusion between the two processes, however further work should involve improved characterization of the glutamate response function combined with synthetic studies to confirm this hypothesis.

In this work we assume the BOLD linewidth change is modelled with a purely Lorentzian character, arising from the assumption of a monoexponential signal decay associated with changes in tissue T2* relaxation^46^. This model is supported by several examples in the literature^4,10,27^ where data from the active portion of the task is Lorentzian linebroadened to match the rest phase and subtracted. These subtraction spectra show very low levels of signal from the primary singlet metabolites (tNAA, tCr and tCho) while maintaining signal from task related metabolites such as lactate and glutamate, supporting the assumption of a Lorentzian lineshape change. A similar procedure with a purely Gaussian character would produce much larger residuals, however we cannot rule out a small non-Lorentzian contribution to the lineshape change and this area warrants further study.

In conclusion, our results support the use of explicit BOLD lineshape correction or modelling for task-based fMRS analysis to reduce fitting bias at field strengths of 3T and above.

## Data Availability

The code required to run all simulations is available online in two repositories. For synthetic data generation and conventional fitting: https://github.com/martin3141/fmrs_bold_bias_paper and for the dynamic fitting approaches: https://github.com/wtclarke/fmrs_bold_dynamic_fitting (commit 39ac814e78a74e47b58510144ab57a5e11f85dcb).

## Acknowledgments

WTC and SF are funded by Wellcome [225924/Z/22/Z]. The Centre for Integrative Neuroimaging was supported by core funding from the Wellcome Trust (203139/Z/16/Z and 203139/A/16/Z).

This research was funded in whole, or in part, by the Wellcome Trust [Grant number 225924/Z/22/Z]. For the purpose of open access, the author has applied a CC BY public copyright licence to any Author Accepted Manuscript version arising from this submission.

## References

1. Mullins PG. Towards a theory of functional magnetic resonance spectroscopy (fMRS): A meta-analysis and discussion of using MRS to measure changes in neurotransmitters in real time. Scand J Psychol. 2018;59(1):91–103. doi:10.1111/sjop.12411

2. Stanley JA, Raz N. Functional Magnetic Resonance Spectroscopy: The “New” MRS for Cognitive Neuroscience and Psychiatry Research. Front Psychiatry. 2018;9. Accessed March 7, 2024. https://www.frontiersin.org/journals/psychiatry/articles/10.3389/fpsyt.2018.00076

3. Prichard J, Rothman D, Novotny E, et al. Lactate rise detected by 1H NMR in human visual cortex during physiologic stimulation. Proc Natl Acad Sci. 1991;88(13):5829–5831. doi:10.1073/pnas.88.13.5829

4. Bednařík P, Tkáč I, Giove F, et al. Neurochemical and BOLD responses during neuronal activation measured in the human visual cortex at 7 Tesla. J Cereb Blood Flow Metab Off J Int Soc Cereb Blood Flow Metab. 2015;35(4):601–610. doi:10.1038/jcbfm.2014.233

5. Maddock RJ, Buonocore MH, Lavoie SP, et al. Brain lactate responses during visual stimulation in fasting and hyperglycemic subjects: a proton magnetic resonance spectroscopy study at 1.5 Tesla. Psychiatry Res. 2006;148(1):47–54. doi:10.1016/j.pscychresns.2006.02.004

6. Mangia S, Tkác I, Gruetter R, Van de Moortele PF, Maraviglia B, Uğurbil K. Sustained neuronal activation raises oxidative metabolism to a new steady-state level: evidence from 1H NMR spectroscopy in the human visual cortex. J Cereb Blood Flow Metab Off J Int Soc Cereb Blood Flow Metab. 2007;27(5):1055–1063. doi:10.1038/sj.jcbfm.9600401

7. Mangia S, Tkác I, Logothetis NK, Gruetter R, Van de Moortele PF, Uğurbil K. Dynamics of lactate concentration and blood oxygen level-dependent effect in the human visual cortex during repeated identical stimuli. J Neurosci Res. 2007;85(15):3340–3346. doi:10.1002/jnr.21371

8. Sappey-Marinier D, Calabrese G, Fein G, Hugg JW, Biggins C, Weiner MW. Effect of Photic Stimulation on Human Visual Cortex Lactate and Phosphates Using 1H and 31P Magnetic Resonance Spectroscopy. J Cereb Blood Flow Metab. 1992;12(4):584–592. doi:10.1038/jcbfm.1992.82

9. Schaller B, Mekle R, Xin L, Kunz N, Gruetter R. Net increase of lactate and glutamate concentration in activated human visual cortex detected with magnetic resonance spectroscopy at 7 tesla. J Neurosci Res. 2013;91(8):1076–1083. doi:10.1002/jnr.23194

10. Lin Y, Stephenson MC, Xin L, Napolitano A, Morris PG. Investigating the Metabolic Changes due to Visual Stimulation using Functional Proton Magnetic Resonance Spectroscopy at 7□ T. J Cereb Blood Flow Metab. 2012;32(8):1484–1495. doi:10.1038/jcbfm.2012.33

11. Frahm J, Krüger G, Merboldt KD, Kleinschmidt A. Dynamic uncoupling and recoupling of perfusion and oxidative metabolism during focal brain activation in man. Magn Reson Med. 1996;35(2):143–148. doi:10.1002/mrm.1910350202

12. Mullins PG, Rowland LM, Jung RE, Sibbitt WL. A novel technique to study the brain’s response to pain: Proton magnetic resonance spectroscopy. NeuroImage. 2005;26(2):642–646. doi:10.1016/j.neuroimage.2005.02.001

13. Gussew A, Rzanny R, Erdtel M, et al. Time-resolved functional 1H MR spectroscopic detection of glutamate concentration changes in the brain during acute heat pain stimulation. NeuroImage. 2010;49(2):1895–1902. doi:10.1016/j.neuroimage.2009.09.007

14. Jelen LA, Lythgoe DJ, Jackson JB, Howard MA, Stone JM, Egerton A. Imaging Brain Glx Dynamics in Response to Pressure Pain Stimulation: A 1H-fMRS Study. Front Psychiatry. 2021;12:681419. doi:10.3389/fpsyt.2021.681419

15. Volovyk O, Tal A. Increased Glutamate concentrations during prolonged motor activation as measured using functional Magnetic Resonance Spectroscopy at 3T. NeuroImage. 2020;223:117338. doi:10.1016/j.neuroimage.2020.117338

16. Morelli M, Dudzikowska K, Deelchand DK, et al. Functional magnetic resonance spectroscopy of prolonged motor activation using conventional and spectral GLM analyses. Imaging Neurosci. 2025;3:imag_a_00452. doi:10.1162/imag_a_00452

17. Schaller B, Xin L, O’Brien K, Magill AW, Gruetter R. Are glutamate and lactate increases ubiquitous to physiological activation? A 1H functional MR spectroscopy study during motor activation in human brain at 7Tesla. NeuroImage. 2014;93:138–145. doi:10.1016/j.neuroimage.2014.02.016

18. Apšvalka D, Gadie A, Clemence M, Mullins PG. Event-related dynamics of glutamate and BOLD effects measured using functional magnetic resonance spectroscopy (fMRS) at 3T in a repetition suppression paradigm. NeuroImage. 2015;118:292–300. doi:10.1016/j.neuroimage.2015.06.015

19. Koolschijn RS, Clarke WT, Ip IB, Emir UE, Barron HC. Event-related functional magnetic resonance spectroscopy. NeuroImage. 2023;276:120194. doi:10.1016/j.neuroimage.2023.120194

20. DiNuzzo M, Mangia S, Moraschi M, Mascali D, Hagberg GE, Giove F. Perception is associated with the brain’s metabolic response to sensory stimulation. eLife. 2022;11:e71016. doi:10.7554/eLife.71016

21. Hennig J, Emst Th, Speck O, Deuschl G, Feifel E. Detection of brain activation using oxygenation sensitive functional spectroscopy. Magn Reson Med. 1994;31(1):85–90. doi:10.1002/mrm.1910310115

22. Zhu XH, Chen W. Observed BOLD effects on cerebral metabolite resonances in human visual cortex during visual stimulation: A functional 1H MRS study at 4 T. Magn Reson Med. 2001;46(5):841–847. doi:10.1002/mrm.1267

23. Provencher SW. Estimation of metabolite concentrations from localized in vivo proton NMR spectra. Magn Reson Med. 1993;30(6):672–679. doi:10.1002/mrm.1910300604

24. Marjańska M, Deelchand DK, Kreis R. Results and interpretation of a fitting challenge for magnetic resonance spectroscopy set up by the MRS study group of ISMRM. Magn Reson Med. 2022;87(1):11–32. doi:10.1002/mrm.28942

25. Mangia S, Tkáč I, Gruetter R, et al. Sensitivity of single-voxel 1H-MRS in investigating the metabolism of the activated human visual cortex at 7 T. Magn Reson Imaging. 2006;24(4):343–348. doi:10.1016/j.mri.2005.12.023

26. van der Zwaag W, Francis S, Head K, et al. fMRI at 1.5, 3 and 7 T: Characterising BOLD signal changes. NeuroImage. 2009;47(4):1425–1434. doi:10.1016/j.neuroimage.2009.05.015

27. Boillat Y, Xin L, Van Der Zwaag W, Gruetter R. Metabolite concentration changes associated with positive and negative BOLD responses in the human visual cortex: A functional MRS study at 7 Tesla. J Cereb Blood Flow Metab. 2020;40(3):488–500. doi:10.1177/0271678X19831022

28. Near J, Andersson J, Maron E, et al. Unedited in vivo detection and quantification of γ-aminobutyric acid in the occipital cortex using short-TE MRS at 3 T. NMR Biomed. 2013;26(11):1353–1362. doi:10.1002/nbm.2960

29. Near J, Harris AD, Juchem C, et al. Preprocessing, analysis and quantification in single-voxel magnetic resonance spectroscopy: experts’ consensus recommendations. NMR Biomed. 2021;34(5):e4257. doi:10.1002/nbm.4257

30. Hui SCN, Mikkelsen M, Zöllner HJ, et al. Frequency drift in MR spectroscopy at 3T. NeuroImage. 2021;241:118430. doi:10.1016/j.neuroimage.2021.118430

31. Clarke WT, Ligneul C, Cottaar M, Ip IB, Jbabdi S. Universal dynamic fitting of magnetic resonance spectroscopy. Magn Reson Med. 2024;91(6):2229–2246. doi:10.1002/mrm.30001

32. Tal A. The future is 2D: spectral-temporal fitting of dynamic MRS data provides exponential gains in precision over conventional approaches. Magn Reson Med. 2023;89(2):499–507. doi:10.1002/mrm.29456

33. Chong DGQ, Kreis R, Bolliger CS, Boesch C, Slotboom J. Two-dimensional linear-combination model fitting of magnetic resonance spectra to define the macromolecule baseline using FiTAID, a Fitting Tool for Arrays of Interrelated Datasets. Magn Reson Mater Phys Biol Med. 2011;24(3):147–164. doi:10.1007/s10334-011-0246-y

34. Govindaraju V, Young K, Maudsley AA. Proton NMR chemical shifts and coupling constants for brain metabolites. NMR Biomed. 2000;13(3):129–153. doi:10.1002/1099-1492(200005)13:3%3C129::aid-nbm619%3E3.0.co;2-v

35. Govind V, Young K, Maudsley AA. Corrigendum: proton NMR chemical shifts and coupling constants for brain metabolites. Govindaraju V, Young K, Maudsley AA, NMR Biomed. 2000; 13: 129-153. NMR Biomed. 2015;28(7):923–924. doi:10.1002/nbm.3336

36. Wilson M, Reynolds G, Kauppinen RA, Arvanitis TN, Peet AC. A constrained least-squares approach to the automated quantitation of in vivo 1H magnetic resonance spectroscopy data. Magn Reson Med. 2011;65(1):1–12. doi:10.1002/mrm.22579

37. Oz G, Tkáč I. Short-echo, single-shot, full-intensity proton magnetic resonance spectroscopy for neurochemical profiling at 4 T: validation in the cerebellum and brainstem. Magn Reson Med. 2011;65(4):901–910. doi:10.1002/mrm.22708

38. Lindquist MA, Loh JM, Atlas LY, Wager TD. Modeling the Hemodynamic Response Function in fMRI: Efficiency, Bias and Mis-modeling. Neuroimage. 2009;45(1 Suppl):S187–S198. doi:10.1016/j.neuroimage.2008.10.065

39. Clarke WT, Bell TK, Emir UE, et al. NIfTI-MRS: A standard data format for magnetic resonance spectroscopy. Magn Reson Med. Published online September 11, 2022. doi:10.1002/mrm.29418

40. Bouchard AE, Wong D, Bogner W, et al. MRS-BIDS, an extension to the Brain Imaging Data Structure for magnetic resonance spectroscopy. Sci Data. 2025;12(1):1384. doi:10.1038/s41597-025-05543-2

41. Wilson M. Robust retrospective frequency and phase correction for single-voxel MR spectroscopy. Magn Reson Med. 2019;81(5):2878–2886. doi:10.1002/mrm.27605

42. Wilson M. Chemical shift and relaxation regularization improve the accuracy of 1H MR spectroscopy analysis. Magn Reson Med. 2025;93(6):2287–2296. doi:10.1002/mrm.30462

43. Wilson M. spant: An R package for magnetic resonance spectroscopy analysis. J Open Source Softw. 2021;6(67):3646. doi:10.21105/joss.03646

44. Clarke WT, Stagg CJ, Jbabdi S. FSL-MRS: An end-to-end spectroscopy analysis package. Magn Reson Med. 2021;85(6):2950–2964. doi:10.1002/mrm.28630

45. Woolrich MW, Behrens TEJ, Beckmann CF, Jenkinson M, Smith SM. Multilevel linear modelling for FMRI group analysis using Bayesian inference. NeuroImage. 2004;21(4):1732–1747. doi:10.1016/j.neuroimage.2003.12.023

46. Speck O, Hennig J. Functional Imaging by I0- and T2* -parameter mapping using multi-image EPI. Magn Reson Med. 1998;40(2):243–248. doi:10.1002/mrm.1910400210

