## appendix a for "The Impact of BOLD Induced Linewidth Modulation on Functional ^1^H MRS Analysis"

Appendix A: Asymptotic derivation of the effect of Lorentzian broadening on spectral fitting

### Problem Summary

Here we asymptotically derive the impact of Lorentzian broadening (apodisation) on the bias and variance of model parameters through both ordinary least squares (OLS) and generalised least squares (GLS) spectral fitting. Below, we define the cost function for both least squares methods, the asymptotic expansion around the true parameter values, and finally the leading-order bias and variance under this asymptotic expansion. We then present a simple forward model for spectral fitting (single Lorentzian peak) as well as analytical expressions for the Jacobian and Hessian. Finally, we analytically derive the noise covariance matrix in both the time and spectral domains when apodised by a spectral line-broadening window. Validation of the asymptotic solutions via comparison to Monte-Carlo simulations is present in the supplementary materials (see Figures S3-S5).

### The Model and Cost Function

Let the observed complex data be $\boldsymbol{Z}$, generated by the true parameters $\boldsymbol{\theta}_{0}$,

$$\boldsymbol{Z}=M(\boldsymbol{\theta}_{0})+\boldsymbol{\epsilon},$$

$M$ is a general forward model and the noise is complex circular Gaussian, $\boldsymbol{\epsilon}\sim\mathcal{N}_{\mathcal{C}}(\mathbf{0},\boldsymbol{\Sigma})$.

We seek the estimate $\hat{\boldsymbol{\theta}}=\boldsymbol{\theta}_{0}+\Delta\boldsymbol{\theta}$ that minimizes the generalized cost function with a weighting matrix $\boldsymbol{\Omega}$:

$$\begin{aligned} C(\boldsymbol{\theta})=(\boldsymbol{Z}-M\left( \boldsymbol{\theta} \right))^{H}\boldsymbol{\Omega}\left( \boldsymbol{Z}-M\left( \boldsymbol{\theta} \right) \right). \#\left( 1 \right) \end{aligned}$$

For now, we will keep the weighting matrix general, however we note that for ordinary least squares, $\boldsymbol{\Omega}=\boldsymbol{I}$ and for generalized least squares $\boldsymbol{\Omega}=\boldsymbol{\Sigma}^{\boldsymbol{-1}}$**,** where $\boldsymbol{\Sigma}$ is the known noise covariance matrix (derived later). Setting the gradient of Equation (1) with respect to $\boldsymbol{\theta}$ to zero at $\hat{\boldsymbol{\theta}}$ gives,

$$\begin{aligned} \nabla C\left( \hat{\boldsymbol{\theta}} \right)=-2\mathrm{Re}\left[ J(\hat{\boldsymbol{\theta}})^{H}\boldsymbol{\Omega}(\boldsymbol{Z}-M(\hat{\boldsymbol{\theta}})) \right]=\mathbf{0,}\#\left( \mathbf{2} \right) \end{aligned}$$

where $J(\boldsymbol{\theta})=\nabla_{\boldsymbol{\theta}}M(\boldsymbol{\theta})$ is the Jacobian.

### Asymptotic Expansion

We proceed by defining the asymptotic parameter as the typical magnitude of the noise $\epsilon\sim|\boldsymbol{\epsilon}|$. The asymptotic assumption holds when $\epsilon\ll M(\boldsymbol{\theta}_{0})$ such that the impact of the apodised noise may be considered a perturbation on the true parameter values, $\boldsymbol{\theta}_{0}$. It will be convenient to expand $\hat{\boldsymbol{\theta}}$ to second order,

$$\begin{aligned} \hat{\boldsymbol{\theta}}=\boldsymbol{\theta}_{0}+\epsilon\boldsymbol{\theta}_{1}+\epsilon^{2}\boldsymbol{\theta}_{2}\mathcal{+O}\left( \epsilon^{3} \right). \#\left( 3 \right) \end{aligned}$$

Here, $\boldsymbol{\theta}_{i}$ is the $i$-th order contribution to the asymptotic approximation of $\hat{\boldsymbol{\theta}}$. Since $\epsilon$ is small, each consecutive term in this expansion is less impactful on the resulting solution. The total error in the estimate may be recovered via $\Delta\boldsymbol{\theta}=\epsilon\boldsymbol{\theta}_{1}+\epsilon^{2}\boldsymbol{\theta}_{2}\mathcal{+O}\left( \epsilon^{3} \right)$.

Using the expansion of $\hat{\boldsymbol{\theta}}$ in Equation (3), we Taylor expand $M(\hat{\boldsymbol{\theta}})$ and $J(\hat{\boldsymbol{\theta}})$ around the true parameters $\boldsymbol{\theta}_{0}$. Let $J=J(\boldsymbol{\theta}_{0})$ and let $H=\nabla_{\boldsymbol{\theta}}^{2}M(\boldsymbol{\theta}_{0})$ be the Hessian tensor.

The expansions up to second order in $\epsilon$ are:

$$\begin{aligned} M(\hat{\boldsymbol{\theta}})&\approx M(\boldsymbol{\theta}_{0})+\epsilon J\boldsymbol{\theta}_{1}+\epsilon^{2}J\boldsymbol{\theta}_{2}+\frac{1}{2}\epsilon^{2}\boldsymbol{\theta}_{1}^{T}H\boldsymbol{\theta}_{1}, \\ \boldsymbol{Z}-M(\hat{\boldsymbol{\theta}})&\approx\boldsymbol{\epsilon}-\epsilon J\boldsymbol{\theta}_{1}-\epsilon^{2}J\boldsymbol{\theta}_{2}-\frac{1}{2}\epsilon^{2}\boldsymbol{\theta}_{1}^{T}H\boldsymbol{\theta}_{1}, \\ J(\hat{\boldsymbol{\theta}})&\approx J+\epsilon\boldsymbol{\theta}_{1}\cdot H, \end{aligned}$$

where $\cdot$ denotes tensor contraction over the parameter dimension. Substituting these into Equation (2) yields:

$$\begin{aligned} \mathbf{0}=Re\left[ (J+\epsilon\boldsymbol{\theta}_{1}\cdot H)^{H}\boldsymbol{\Omega}\left( \boldsymbol{\epsilon}-\epsilon J\boldsymbol{\theta}_{1}\epsilon^{2}J\boldsymbol{\theta}_{2}-\frac{1}{2}\epsilon^{2}\boldsymbol{\theta}_{1}^{T}H\boldsymbol{\theta}_{1} \right) \right]. \boldsymbol{\#}\left( 4 \right) \end{aligned}$$

### First-Order Error and Variance

We aim to asymptotically predict the bias $\mathbb{E}\left[ \Delta\boldsymbol{\theta} \right]$ and variance $\mathbb{E}\left[ \Delta\boldsymbol{\theta}\Delta\boldsymbol{\theta}^{T} \right]$ of predicted offset due to the apodised noise. To do this, we first consider the first-order terms (linear in $\epsilon$). Extracting the $\mathcal{O(}\epsilon)$ terms from Equation (4) gives:

$$\mathbf{0}=Re\left[ J^{H}\boldsymbol{\Omega}\boldsymbol{\epsilon}-J^{H}\boldsymbol{\Omega}J\boldsymbol{\theta}_{1} \right].$$

Solving for the first-order error:

$$\begin{aligned} \boldsymbol{\theta}_{1}=\left( \mathrm{Re}\left[ J^{H}\boldsymbol{\Omega}J \right] \right)^{-1}\mathrm{Re}\left[ J^{H}\boldsymbol{\Omega}\boldsymbol{\epsilon} \right]=\boldsymbol{M}_{\Omega}\mathrm{Re}\left[ J^{H}\boldsymbol{\Omega}\boldsymbol{\epsilon} \right], \#\left( 5 \right) \end{aligned}$$

where $\boldsymbol{M}_{\Omega}=(Re[J^{H}\boldsymbol{\Omega}J])^{-1}$ is the unweighted projection matrix. Since $\mathbb{E[}\boldsymbol{\epsilon}]=\mathbf{0}$, the predicted first-order bias is zero. The parameter variance is given by

$$\begin{aligned} \boldsymbol{V}_{\Omega}\mathbb{=E}\left[ \boldsymbol{\theta}_{1}\boldsymbol{\theta}_{1}^{T} \right]. \#\left( 6 \right) \end{aligned}$$

Substituting Equation (5) into Equation (6), using the identity

$$\mathbb{E}[Re\{x\}Re\{x{\}}^{T}]=\frac{1}{2}Re\{\mathbb{E[}xx^{H}\mathbb{]+E[}xx^{T}]\},$$

and assuming $Re\{\mathbb{E[}\boldsymbol{\epsilon}\boldsymbol{\epsilon}^{T}]\}=\mathbf{0}$ we obtain

$$\begin{aligned} \boldsymbol{V}_{\Omega}=\boldsymbol{M}_{\Omega}\mathrm{Re}\left[ J^{H}\boldsymbol{\Omega\Sigma\Omega}J \right]\boldsymbol{M}_{\Omega}, \#\left( 7 \right) \end{aligned}$$

where $\boldsymbol{\Sigma}\mathbb{=E[}\boldsymbol{\epsilon}\boldsymbol{\epsilon}^{H}]$ is the noise covariance matrix.

Since the first-order bias is zero (and the first-order variance non-zero), this suggests that the primary impact of the apodised noise on the resulting distribution is to increase in variance. This is confirmed in the later comparison to MC simulations.

### Second-Order Error and Bias

In order to calculate the leading-order impact of the apodised noise on the parameter estimation bias, we must consider the second-order asymptotic problem (terms quadratic in $\epsilon$). Extracting the $\mathcal{O(}\epsilon^{2})$ terms from Equation (4) gives:

$$\mathbf{0}=Re\left[ -J^{H}\boldsymbol{\Omega}J\boldsymbol{\theta}_{2}-\frac{1}{2}J^{H}\boldsymbol{\Omega}\left( \boldsymbol{\theta}_{1}^{T}H\boldsymbol{\theta}_{1} \right)+(\boldsymbol{\theta}_{1}\cdot H)^{H}\boldsymbol{\Omega}\boldsymbol{\epsilon}-(\boldsymbol{\theta}_{1}\cdot H)^{H}\boldsymbol{\Omega}J\boldsymbol{\theta}_{1} \right].$$

Solving for $\boldsymbol{\theta}_{2}$ and taking the expectation $\boldsymbol{b}\mathbb{=E[}\boldsymbol{\theta}_{2}]$ yields the leading-order asymptotic bias,

$$\boldsymbol{b}_{\Omega}=-\frac{1}{2}\boldsymbol{M}_{\Omega}\mathrm{Re}\left[ J^{H}\boldsymbol{\Omega}\mathbb{E}\left[ \boldsymbol{\theta}_{1}^{T}H\boldsymbol{\theta}_{1} \right] \right]+\boldsymbol{M}_{\Omega}\mathrm{Re}\left[ \mathbb{E}\left[ (\boldsymbol{\theta}_{1}\cdot H)^{H}\boldsymbol{\Omega}\boldsymbol{\epsilon} \right]\mathbb{-E}\left[ (\boldsymbol{\theta}_{1}\cdot H)^{H}\boldsymbol{\Omega}J\boldsymbol{\theta}_{1} \right] \right].$$

It may be shown that this reduces to

$$\begin{aligned} \boldsymbol{b}_{\Omega}=-\frac{1}{2}\boldsymbol{M}_{\Omega}\mathrm{Re}\left[ J^{H}\boldsymbol{\Omega}H:\boldsymbol{V}_{\Omega} \right]+M_{\Omega}\mathrm{Re}\left[ \frac{1}{2}\boldsymbol{\Omega\Sigma\Omega}J\boldsymbol{M}_{\Omega}-\boldsymbol{\Omega}J\boldsymbol{V}_{\Omega} \right], \#\left( 8 \right) \end{aligned}$$

where $:$ is a contraction between tensors.

### OLS vs GLS

For OLS (not whitened), we substitute $\boldsymbol{\Omega}=\boldsymbol{I}$ into the general equations for the variance and bias, Equations (7-8). Simplifying, we obtain the leading order equations

$$\begin{aligned} \begin{aligned} \boldsymbol{V}_{U}&=\frac{1}{2}\boldsymbol{M}_{U}\mathrm{Re}\left[ J^{H}\boldsymbol{\Sigma}J \right]\boldsymbol{M}_{U}, \text{where} \boldsymbol{M}_{U}=\left( \mathrm{Re}\left[ J^{H}J \right] \right)^{-1}, \# \\ \boldsymbol{b}_{U}&=-\frac{1}{2}\boldsymbol{M}_{U}\mathrm{Re}\left[ J^{H}H:\boldsymbol{V}_{U} \right]+\boldsymbol{M}_{U}\mathrm{Re}\left[ \frac{1}{2}\boldsymbol{\Sigma}J\boldsymbol{M}_{U}-J\boldsymbol{V}_{U} \right]. \end{aligned}\#\left( 9 \right) \end{aligned}$$

Conversely, for GLS (whitened residual), we substitute $\boldsymbol{\Omega}=\boldsymbol{\Sigma}^{-1}$ into Equations (7-8) to obtain

$$\begin{aligned} \begin{aligned} \boldsymbol{V}_{W}&=\frac{1}{2}\boldsymbol{M}_{\boldsymbol{W}}=\frac{1}{2}\left( \mathrm{Re}\left[ J^{H}\boldsymbol{\Sigma}^{-1}J \right] \right)^{-1}, \\ \boldsymbol{b}_{W}&=-\frac{1}{2}\boldsymbol{M}_{W}\mathrm{Re}\left[ J^{H}\boldsymbol{\Sigma}^{-1}H:\boldsymbol{V}_{W} \right]. \end{aligned}\#\left( 10 \right) \end{aligned}$$

We note that the last term in Equation (8) cancels for GLS.

### Lorentzian model

We now present a simple forward model to test the impacts of apodisation, a single Lorentzian peak defined by a concentration, shift, and line-width. The Jacobian and Hessian may be substituted into the above expressions for the bias and variance, Equations (9-10).

#### Definitions and Base Terms

We define the parameter vector as $\boldsymbol{x}=[c,s,\gamma]^{\top}$, where $c$ is the concentration, $s$ is the frequency shift, and $\gamma$ is the line-broadening.

Let $b(t)$ represent the original basis function in the time domain. The parametrised time-domain basis signal $m(t)$ is given by:

$$m(t)=exp\left( -(is+\gamma)t \right)b(t)$$

To simplify the notation for the frequency-domain signal and its derivatives, let the Fourier transform ($\mathcal{F}$) of the time-weighted signals be defined as $F_{k}$:

$$\begin{aligned} F_{0}&\mathcal{=F\{}m(t)\} \\ F_{1}&\mathcal{=F\{}t\cdot m(t)\} \\ F_{2}&\mathcal{=F\{}t^{2}\cdot m(t)\} \end{aligned}$$

#### 1. Forward Model ($S$)

The frequency-domain forward model $S$ is the concentration scaled by the zeroth-order Fourier transform of the basis:

$$S(c,s,\gamma)=c\cdot F_{0}$$

#### 2. Jacobian ($J$)

The Jacobian is the matrix of first-order partial derivatives of the forward model with respect to each parameter. For a single spectral point, it is a $1\times3$ row vector:

$$J=\left[ \begin{matrix} \frac{\partial S}{\partial c} & \frac{\partial S}{\partial s} & \frac{\partial S}{\partial\gamma} \end{matrix} \right]$$

Evaluating the partial derivatives of $S$ yields the final Jacobian vector:

$$J=\left[ \begin{matrix} F_{0} & -icF_{1} & -cF_{1} \end{matrix} \right]$$

#### 3. Hessian ($H$)

The Hessian is the $3\times3$ symmetric matrix of second-order partial derivatives:

$$H=\left[ \begin{matrix} \frac{\partial^{2}S}{\partial c^{2}} & \frac{\partial^{2}S}{\partial c\partial s} & \frac{\partial^{2}S}{\partial c\partial\gamma} \\ \frac{\partial^{2}S}{\partial s\partial c} & \frac{\partial^{2}S}{\partial s^{2}} & \frac{\partial^{2}S}{\partial s\partial\gamma} \\ \frac{\partial^{2}S}{\partial\gamma\partial c} & \frac{\partial^{2}S}{\partial\gamma\partial s} & \frac{\partial^{2}S}{\partial\gamma^{2}} \end{matrix} \right]$$

Calculating the individual second-order derivatives and populating the symmetric matrix gives:

$$H=\left[ \begin{matrix} 0 & -iF_{1} & -F_{1} \\ -iF_{1} & -cF_{2} & icF_{2} \\ -F_{1} & icF_{2} & cF_{2} \end{matrix} \right]$$

### Spectral Noise Covariance in MRS after Lorentzian Broadening

#### Time domain

We first define complex white Gaussian noise with $N$ points in the time domain,

$$\begin{aligned} \mathbb{E}\left[ \nu_{n} \right]=0, \text{such that} \mathbb{E}\left[ \nu_{n}\nu_{m}^{*} \right]=\sigma_{t}^{2} \delta_{nm}, \#\left( 11 \right) \end{aligned}$$

$\sigma_{t}^{2}$ is the time-domain variance. We define the apodised (filtered) time-domain noise as

$$\nu_{n}^{'}=\nu_{n}W_{n},$$

where $W_{n}$ is the purely real, time-domain filter (window).

#### Frequency domain

In the frequency (or spectral) domain, the apodised (filtered) noise is the discrete Fourier Transform of $\nu_{n}^{'}$,

$$\tilde{\nu}_{k}=\sum_{n=0}^{N-1} W_{n}\nu_{n} e^{-i2\pi kn/N}.$$

#### Definition of Covariance

The covariance of the apodised noise in the spectral domain is then

$$Cov(\tilde{\nu}_{k},\tilde{\nu}_{\mathcal{l}}\mathbb{)=E}\left[ \left( \sum_{n=0}^{N-1} W_{n}\nu_{n}e^{-i2\pi kn/N} \right)\left( \sum_{m=0}^{N-1} W_{m}\nu_{m}e^{i2\pi\mathcal{l}m/N} \right) \right].$$

Exploiting the linearity of the summation and expectation operators, and using Equation (11), we find

$$Cov(\tilde{\nu}_{k},\tilde{\nu}_{\mathcal{l}})=\sum_{n=0}^{N-1} \sigma_{t}^{2} W_{n}^{2} e^{-i2\pi n(k-\mathcal{l})/N}$$

such that the covariance is the scaled Fourier transform of the square of the time-domain window,

$$Cov(\tilde{\nu}_{k},\tilde{\nu}_{\mathcal{l}})=\sigma_{t}^{2}\times DFT\{W_{n}^{2}\} \text{at frequency index }(k-\mathcal{l}),$$

#### Derivation of the Spectral Noise Covariance for a line-broadening window

Consider the discrete-time window

$$W_{n}=e^{-\gamma n}, n=0,1,\ldots,N-1,$$

where $\gamma>0$, is the Lorentzian line-broadening to be applied.

#### DFT Definition

Applying an $N$-point discrete Fourier transform (DFT) to $W_{n}^{2}$ we have

$$X_{k}=\sum_{n=0}^{N-1} e^{-2\gamma n}e^{-i\frac{2\pi}{N}kn}$$

which may be expressed in closed form as the finite geometric series,

$$X_{k}=\frac{1-e^{-2\gamma N}}{1-e^{-2\gamma}e^{-i\frac{2\pi}{N}k}}.$$

The final covariance matrix can be written as a circulant matrix substituting in the expression $X_{k}$

$$\left( \begin{matrix} X_{N/2} & X_{N/2+1} & X_{N/2+2} & \cdots& X_{N/2-1} \\ X_{N/2-1} & X_{N/2} & X_{N/2+1} & \cdots& X_{N/2-2} \\ X_{N/2-2} & X_{N/2-1} & X_{N/2} & \cdots& X_{N/2-3} \\ \vdots& \vdots& \vdots& \ddots& \vdots\\ X_{N/2+1} & X_{N/2+2} & X_{N/2+3} & \cdots& X_{N/2} \end{matrix} \right).$$
