## supplemental materials for "The Impact of BOLD Induced Linewidth Modulation on Functional ^1^H MRS Analysis"

**Supplementary Materials**

| Basis signal | Concentration (mM) |
| --- | --- |
| Alanine | 0.00 |
| Ascorbate | 0.96 |
| Aspartate | 3.58 |
| Creatine | 4.22 |
| Gamma-aminobutyric acid | 1.03 |
| Glucose | 0.62 |
| Glutamine | 2.79 |
| Glycine | 0.00 |
| Glutathione | 1.09 |
| Glutamate | 8.59 |
| Glycerophosphorylcholine | 0.54 |
| myo-Inositol | 6.08 |
| Lactate | 1.01 |
| N-acetylaspartate | 11.9 |
| N-acetylaspartylglutamate | 1.32 |
| Phosphocholine | 0.40 |
| Phosphocreatine | 3.34 |
| Phosphoethanolamine | 0.93 |
| scyllo-Inositol | 0.27 |
| Taurine | 1.27 |
| Lip09 | 0.00 |
| Lip13a | 0.00 |
| Lip13b | 0.00 |
| Lip20 | 0.00 |
| MM09 | 4.00 |
| MM12 | 4.00 |
| MM14 | 4.00 |
| MM17 | 4.00 |
| MM20 | 4.00 |

Table S1. Molecular signals and associated concentrations used to generate synthetic MRS data.

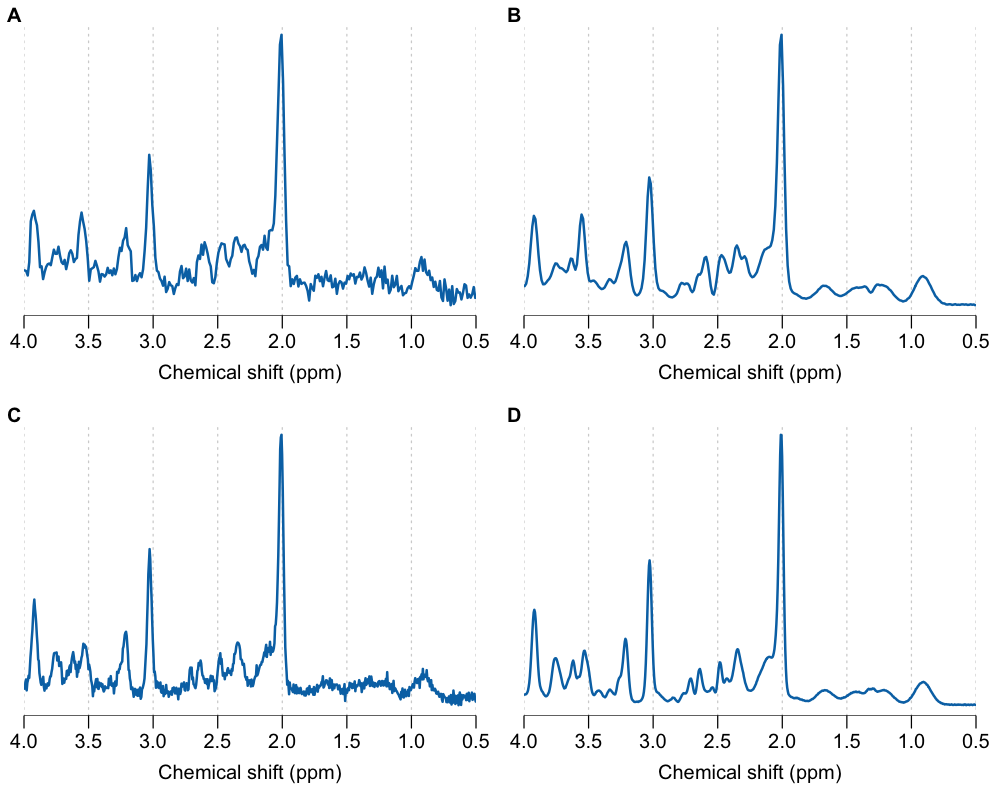

Figure S1. Representative synthetic spectra for a single subject A) single shot 3 T; B) dynamic average (N=640) 3 T; C) single shot 7 T and D) dynamic average (N=320) 7 T.

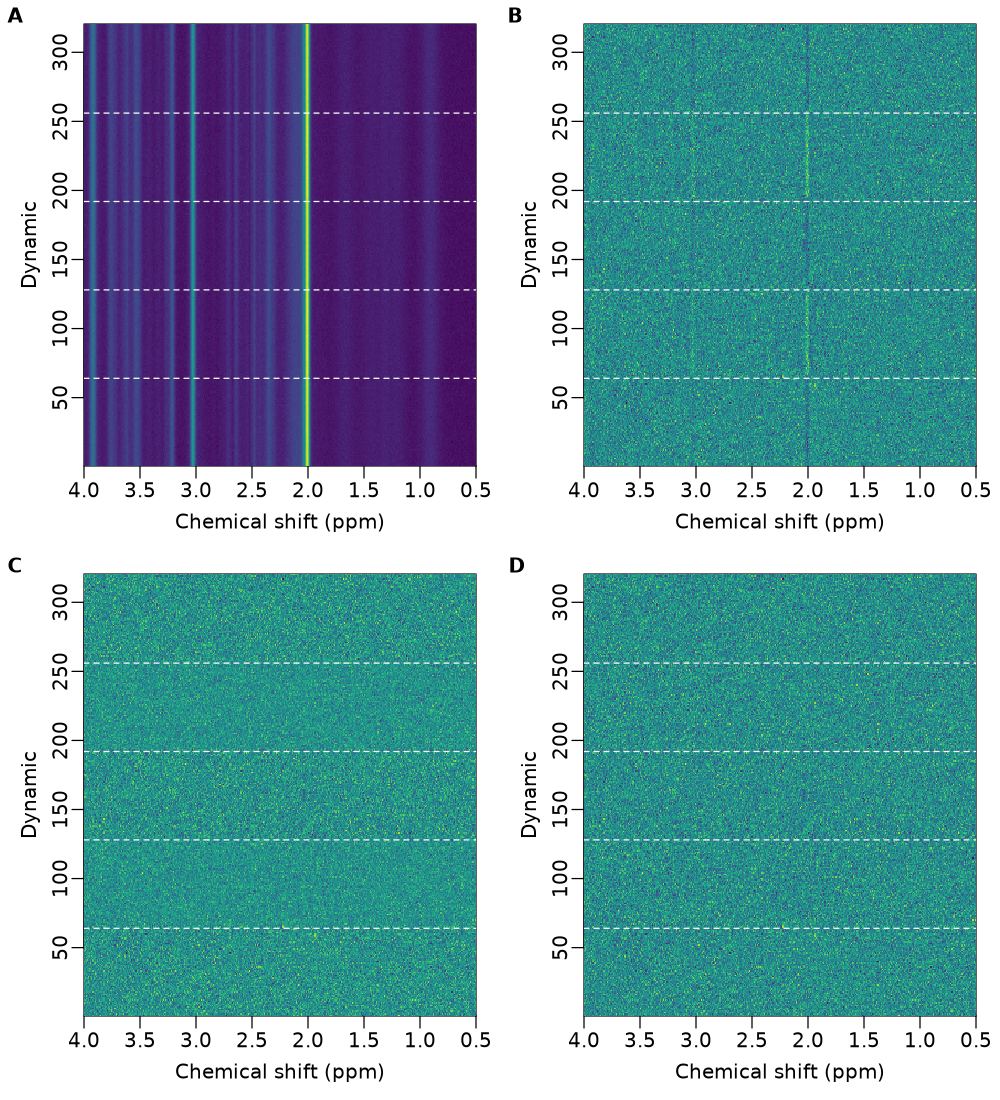

Figure S2. Spectrograms illustrating the Lorentzian lineshape and noise matching preprocessing step with dashed horizontal lines representing the boundaries between the REST - TASK - REST - TASK - REST states. A) unprocessed spectra. B) Unprocessed spectra with the dynamic mean spectrum subtracted. C) Lineshape matched spectra with the dynamic mean spectrum subtracted. D) Lineshape and noise matched spectra with the dynamic mean spectrum subtracted. Data was synthesised with the 7 T parameters listed in the main manuscript.

### Validation of the asymptotic solution via Monte-Carlo simulation

We now validate the asymptotic expressions for the OLS and GLS bias and variance, Equations (9-10), to Monte-Carlo simulations using the forward model and covariance matrix defined in Appendix A. The Monte-Carlo simulations used three noise levels, seven apodisation filters (0 to 20 Hz width), and 20,000 repetitions per condition. Example spectra for each condition are shown in Figure S3. An example comparison of the MC distribution and analytical prediction is presented in Figure S4 with a noise standard deviation of one and an apodisation of 20Hz. The analytical prediction captures the observed MC distribution well, with the bias small in comparison to the variance. The effect of whitening is also clear with the variance of the GLS distribution considerably smaller. To quantify the performance of the analytical predictions, in Figure S4 we compare the NRMSE (%) for both the OLS and GLS optimization approaches over a range of noise and apodisation magnitudes. The analytical predictions are shown to strongly agree with the MC simulations over the range of tested noise and apodisation magnitudes. Unsurprisingly, the largest discrepancy is observed when the noise and apodisation are largest, which we attribute to the asymptotic error in the analytical solutions. We anticipate this discrepancy could be reduced by going to higher orders in the asymptotic solution. The effect of whitening is again clear here, with the NRMSE increasing with apodisation for the OLS solution, but remaining fixed for the GLS solution.

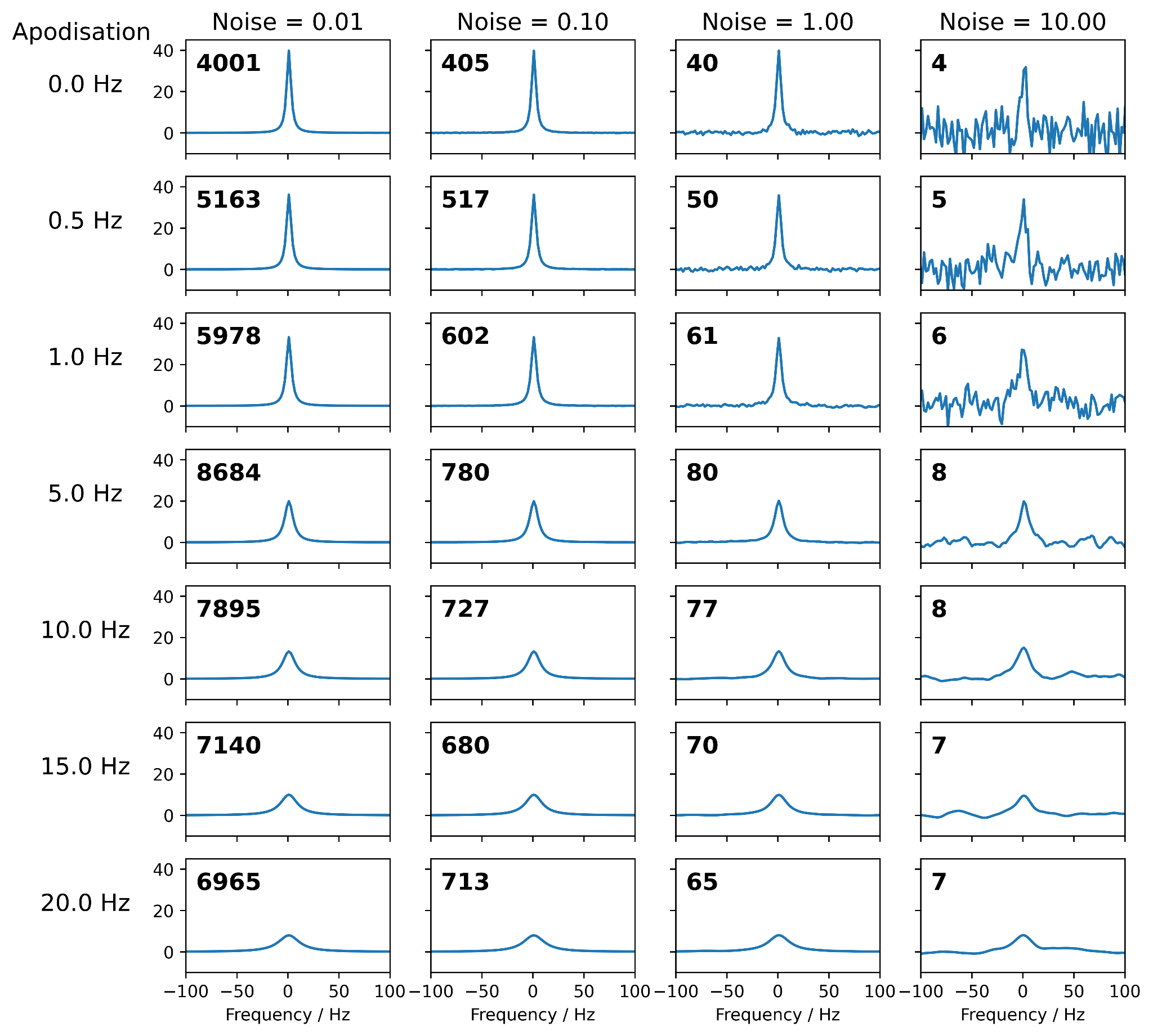

Figure S3. Example single peak spectra from each condition examined for spectral fitting bias after apodisation. The matched filter SNR for each case is printed in the top left. The single peak is lorentzian (FWHM = 5 Hz) and on-resonance (centred at 0 Hz).

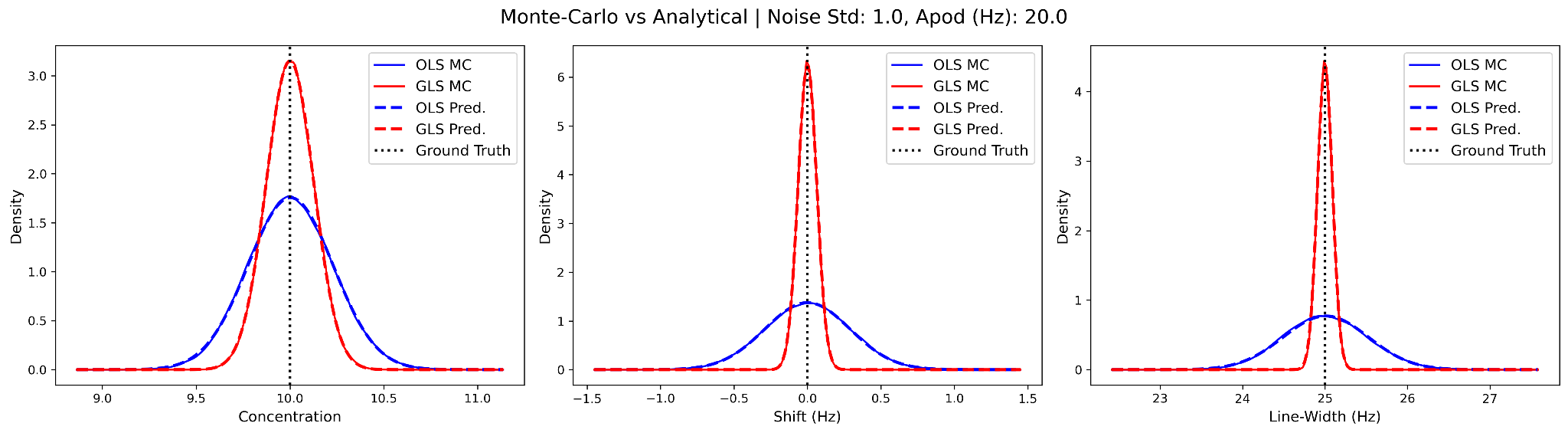

Figure S4: Comparison of concentration, shift, and line-width distributions between the Monte-Carlo (MC) and analytical predictions for both OLS and GLS optimisations with a noise standard deviation of one and an apodisation of 20 Hz.

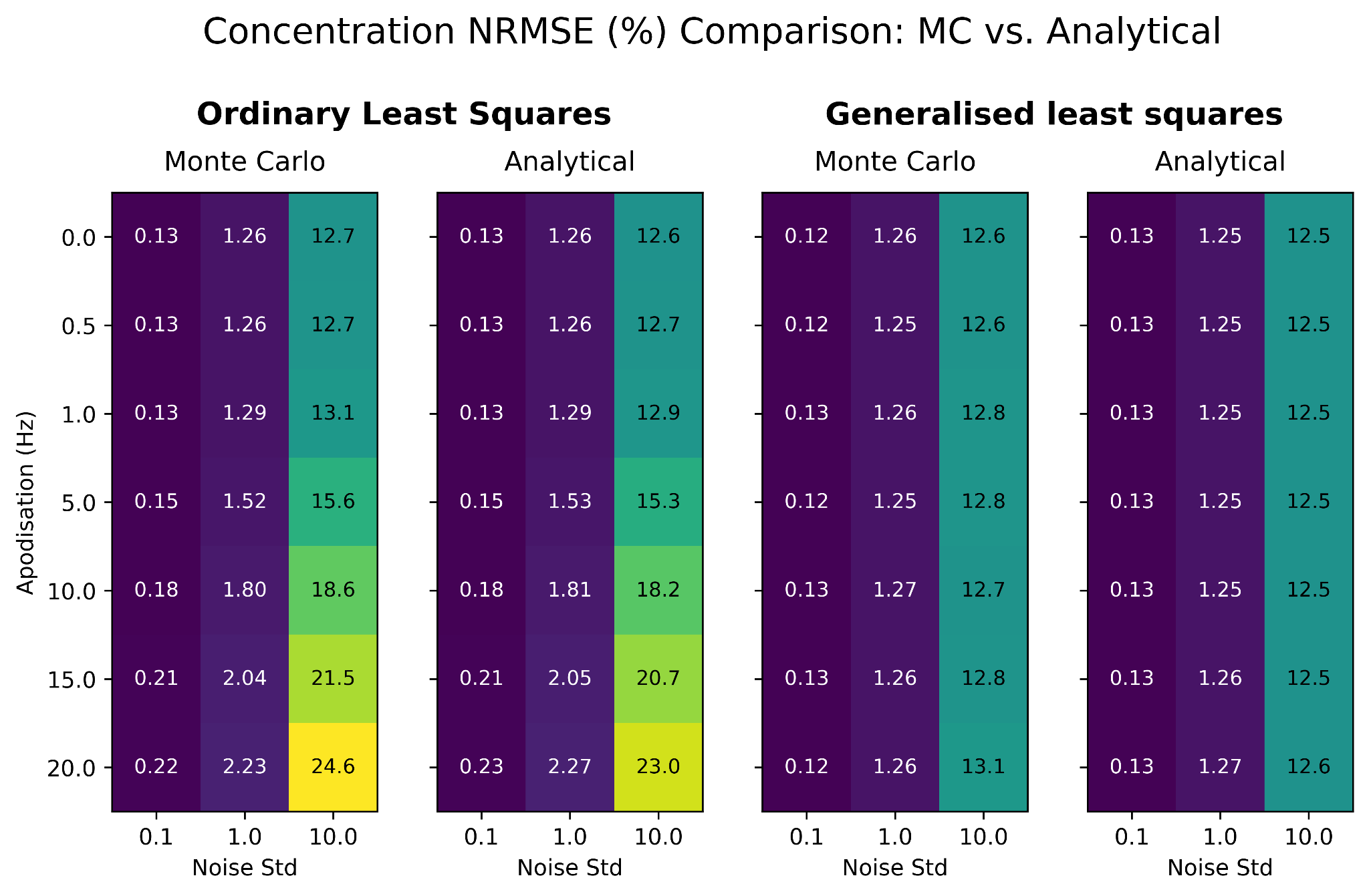

Figure S5: Comparison of the NRMSE (%) between the Monte-Carlo and analytical predictions for both OLS and GLS optimisations over a range of noise standard deviations and apodisation magnitudes.

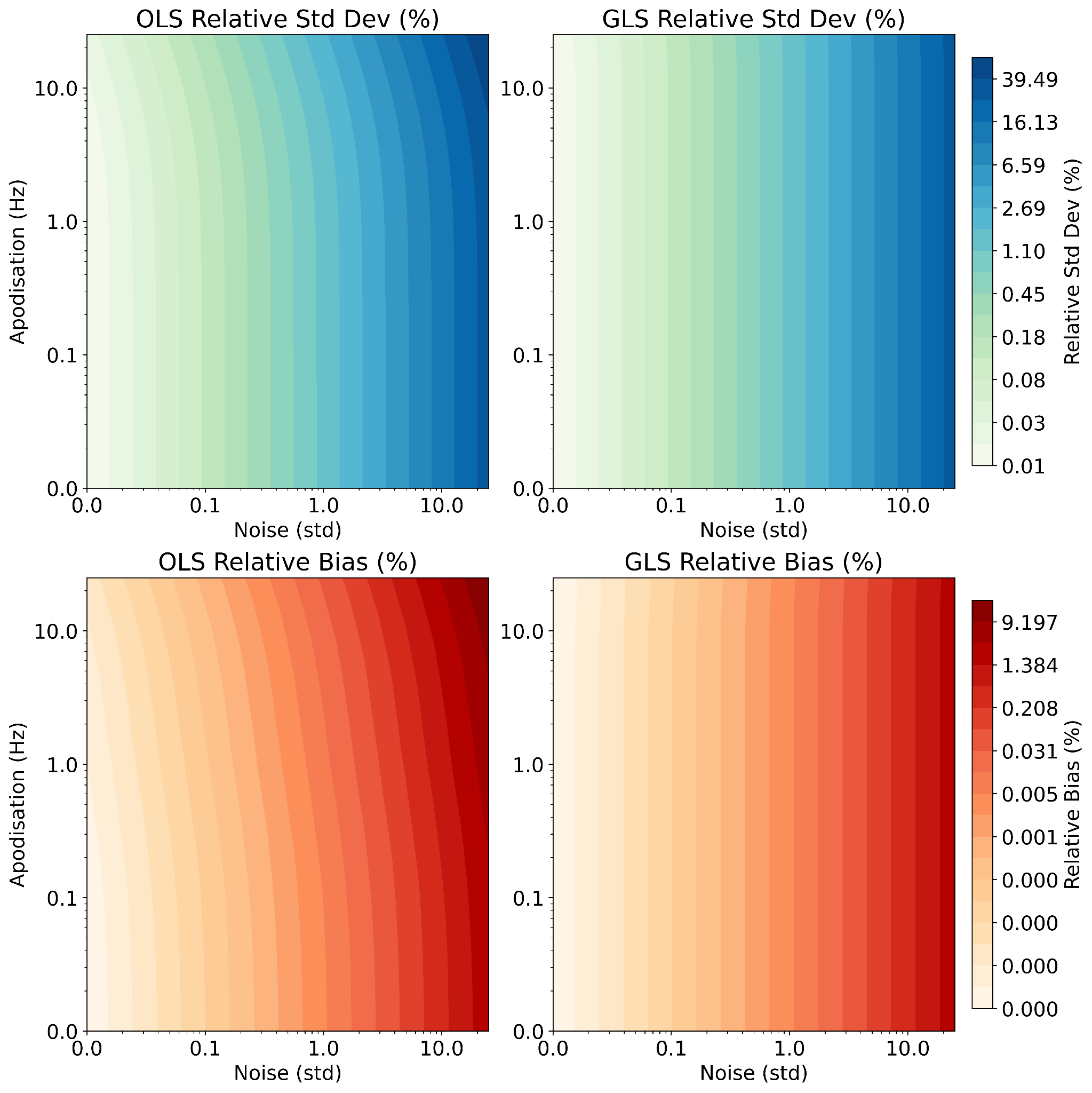
Figure S6: The fitted concentration standard deviation (top) and bias (bottom) predicted using the OLS (left) and GLS (right) asymptotic solutions as a percentage of the ground truth value. Results shown over a range of noise and apodisation magnitudes. Contours and axes both use a log scale for visualisation.

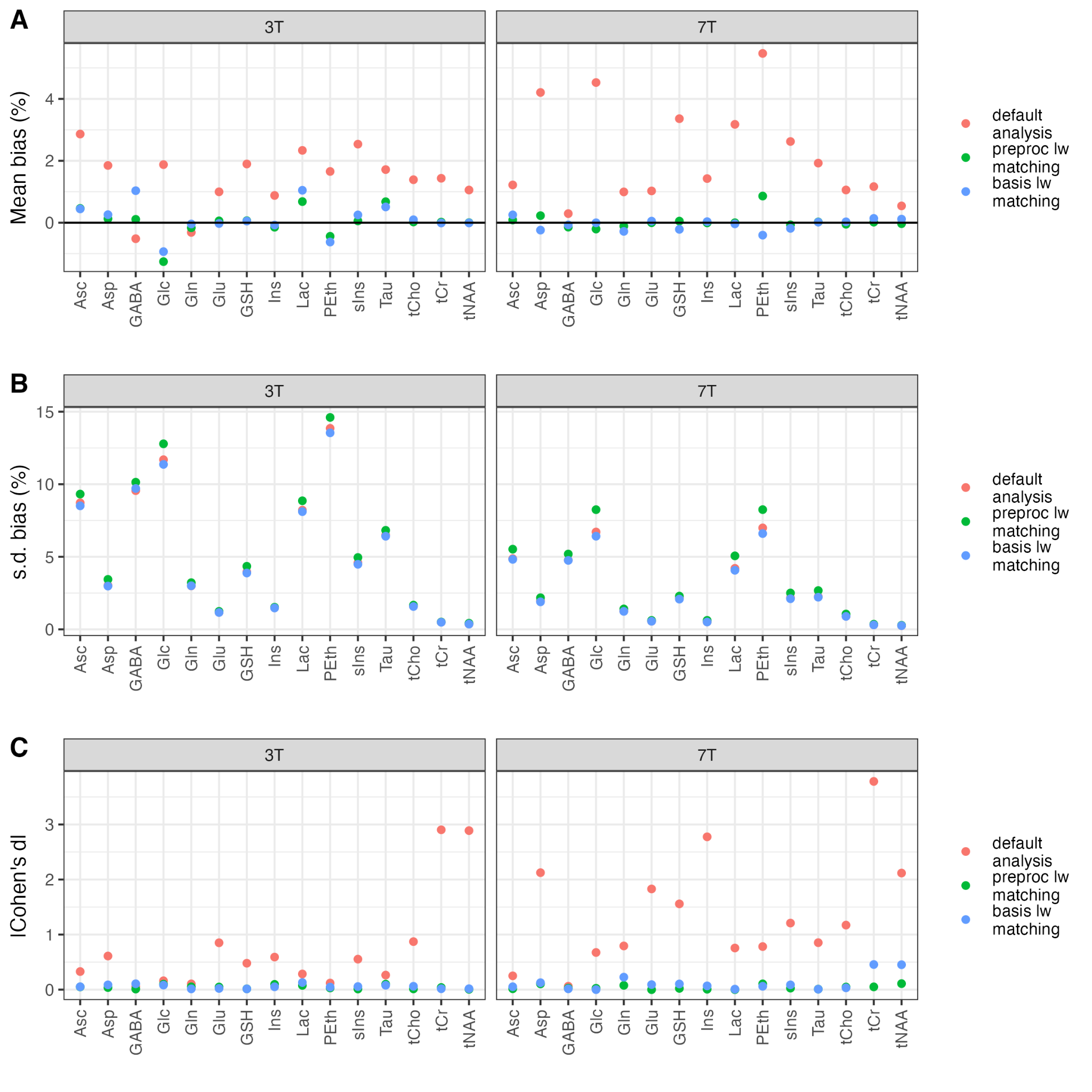

Figure S7. BOLD lineshape induced bias for commonly measured metabolites at 3T and 7T. The LCModel analysis method was used to analyse 5 dynamic blocks (REST - TASK - REST - TASK - REST) and three analysis approaches were compared. Bias was calculated by evaluating the mean of the TASK blocks as a percentage change from the mean of the REST blocks on a participant by participant basis.

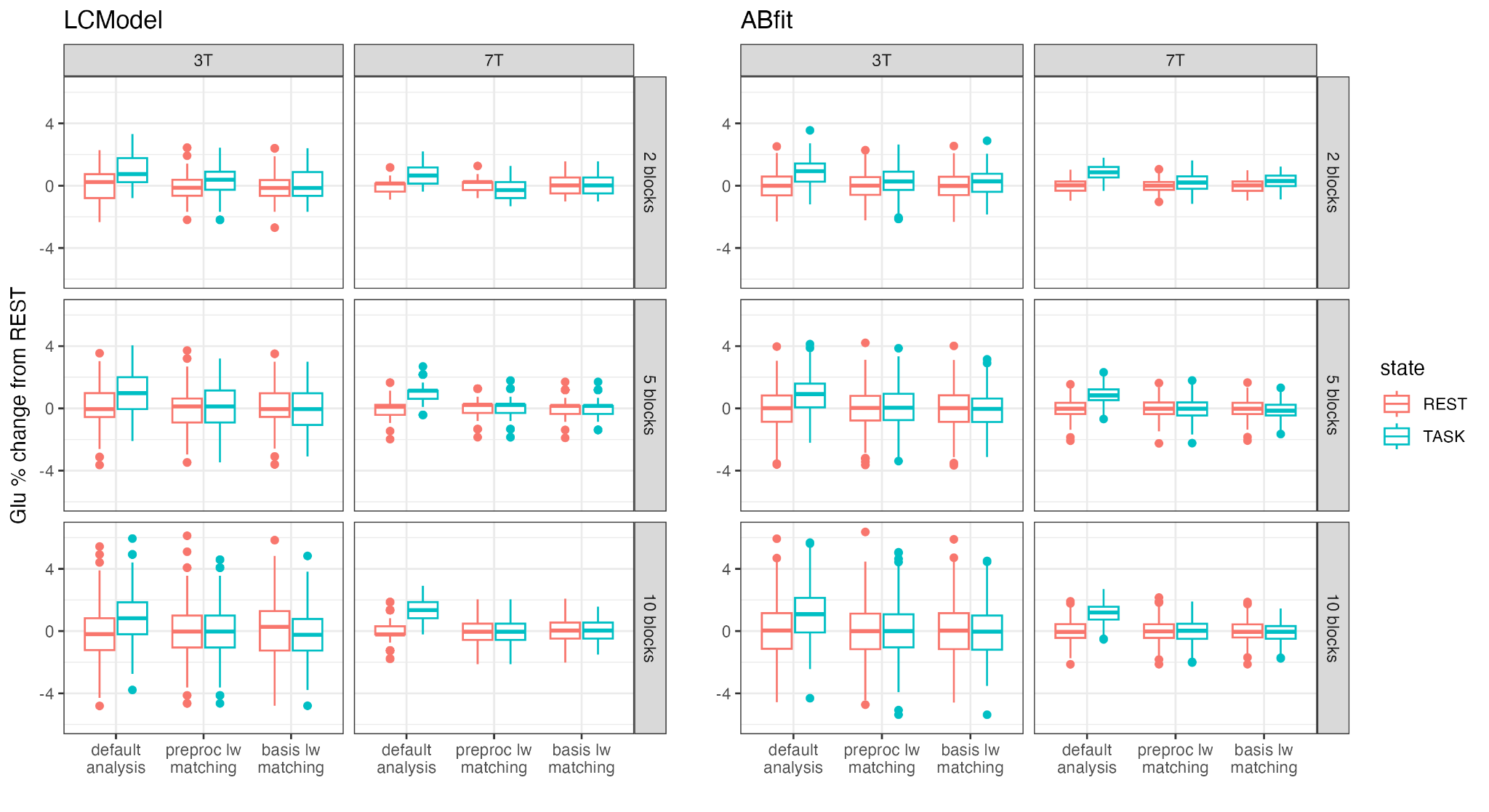

Figure S8. Box and Whisker plots of the percentage change in glutamate level between the REST and TASK conditions. Field strength, number of dynamic blocks, fitting method and BOLD linewidth correction approaches are shown for comparison.
